# *VC1* and the production of vicine and convicine in the genus *Vicia*

**DOI:** 10.64898/2026.07.09.737524

**Authors:** L.L. Vottonen, Wei Chang, Marjo Pöysä, Anna-Maija Lampi, Jaakko Tanskanen, A.H. Schulman, F.L. Stoddard

## Abstract

Many *Vicia* species contain vicine and convicine (VC), which limit the use of faba bean and some vetches in food and feed. The first step in VC biosynthesis in *V. faba* is shared with the riboflavin pathway and attributed to *VC1*, a member of the *ribAB* family. Since riboflavin is ubiquitous to life, we examined the distribution of VC production in genus *Vicia*.

Three accessions of each of 33 *Vicia* species were grown in glasshouse conditions to provide fresh seeds for VC analysis and leaves for DNA analysis. PCR was used to amplify fragments of the *VC1*/*ribAB* gene for sequencing, and these sequences were used to create a phylogenetic tree. *COX1* and *ITS2* sequences were used for examining the nucleotide diversity in the subgenera.

VC and DNA sequences consistent with *VC1* were found only in members of subgenus *Vicia*. In *V. lathyroides, VC1* was present but no VC was detected. There was less sequence diversity in *VC1/ribAB* sequences of subgenus *Cracca* than in those of subgenus *Vicia*, suggesting that *ribAB* remained under stricter purifying selection than *VC1. VC1* is confirmed as a prerequisite for the presence of VC, and the gene and its products are restricted to subgenus *Vicia*.

**Highlight:** The favism-causing factors of vetches and faba bean, vicine and convicine, depend on the presence of the VC1 variant of the ribAB gene, which is found in only one subgenus.

## Introduction

The genus *Vicia* (Fabaceae) comprises between 150 (Kupicha, 1976) and 210 (Hanelt and Mettin, 1989) species. Of these, more than 40 are cultivated (Hanelt and Mettin, 1989), the most important being the faba bean (*Vicia faba* L.). It is grown for human and animal consumption and is one of the oldest domesticated crops. Total world production of faba bean was 6.3 million tonnes in 2024, having grown by 2.5 million tonnes since 2000 (FAOSTAT, 2026). Faba bean is an excellent source of protein, with a protein content of 27–34% of the seed dry matter, and can be grown in temperate climates around the globe (Duc, 1997). Other species in the genus are known as vetches, and those in agricultural use include common vetch (*V. sativa* L.), bitter vetch (*V. ervilia* L.), and hairy vetch (*V. villosa* L.). In the past, these have been used as human food, but now they are primarily used as animal feed and green manure, their use declining with the advent of modern agricultural practices (Van De Wouw *et al*., 2001).

There are many antinutritional factors in vetches, such as tannins, trypsin inhibitors, phenolics (Salehi *et al*., 2021) and, in some, vicine and convicine (VC). VC have been reported only in the genus *Vicia* and in bitter melon (*Momordica charantia* L. (Cucurbitaceae)). These substances are pyrimidine glucosides, which hydrolyze into divicine and isouramil during digestion. The aglycones are powerful oxidants that are harmful to humans who have a genetic mutation conferring a glucose-6-phosphate dehydrogenase (G6PD) deficiency. If a person with G6PD deficiency ingests vicine or convicine, it can cause favism, an acute haemolytic anemia. In the worst cases, favism entails destruction of even 80% of the red blood cells and can be fatal, but most cases are milder (Crépon *et al*., 2010). VC can have negative effects when present in animal feeds; the most affected are chickens, where VC cause decreases in egg size and quality from laying hens and increase mortality when faba bean is included at only 5% of the diet (Koivunen *et al*., 2014). Other monogastric animals such as pigs are also affected by VC, but not as strongly as chickens (Crépon *et al*., 2010). Unlike some other antinutritional factors in legumes that remain in the seedcoat, VC cannot be removed easily in food or feed processing, as they are synthesized in the developing seed coat and stored in the cotyledons and embryo. In wild-type faba bean, VC comprise about 1% of the seed dry matter. The amount varies between cultivars, with low VC cultivars having 5–10% of the wild-type VC (Khamassi *et al*., 2013). The low VC cultivars do not cause problems to humans or animals (Gallo *et al*., 2018).

The biosynthesis of VC was first thought to be derived from the orotic pathway (Brown and Roberts, 1972), but it has recently been shown that VC are products of an overflow mechanism from the riboflavin (vitamin B2) biosynthetic pathway (Björnsdotter *et al*., 2021). The branch point between the overflow to VC and the series of reactions leading to riboflavin is at the step catalyzed by the enzyme encoded by the *ribAB* gene. In faba bean, there are several copies of *ribAB*. One was linked to VC production by the effect of a mutation in its coding sequence and named *VC1*, with the mutant allele *vc1* associated with lower VC levels in faba bean (Björnsdotter *et al*., 2021). Another *ribAB* copy has been linked to the remaining VC production in homozygous *vc1* faba bean plants and named *VC2* (Ugwuanyi *et al*., 2024, Preprint).

During riboflavin synthesis, highly reactive intermediates are formed, and it has been proposed that converting these into VC stops the intermediates from building up in the cell and damaging it (Frelin *et al*., 2015). However, although riboflavin is produced ubiquitously in plants, bacteria and fungi, only in certain *Vicia* species (and in the phylogenetically remote bitter melon, *Momordica charantia* L.), is VC also synthesized. VC production also occurs during germination of the seed and in the roots. The *VC1* gene is connected to VC production in the seed coat, whereas other copies of *ribAB* are associated with riboflavin production in other tissues (Björnsdotter *et al*., 2021).

Although VC has been found in faba bean and other *Vicia* species, its prevalence has not been clear, nor how its occurrence corresponds to the phylogeny of the genus. Hence, we set out to examine the presence of VC in a range of *Vicia* species, chosen to represent the currently accepted subgenera, and to associate the results with the sequence of the *ribAB* genes in each species. Our hypothesis was that VC synthesis arose at a branch point in the phylogeny of the genus leading to a clade containing *V. faba*, that this knowledge would provide a handle on the evolution of the trait, and thereby, how a future breeding strategy might eliminate VC in *V. faba*.

## Materials and Methods

### Germplasm

A set of 24 *Vicia* species was chosen to represent the broad diversity of the genus according to Leht’s (2009) phylogeny. Three accessions per species, from as environmentally diverse sources as possible, were selected. In some cases, three accessions were not available. Seeds for 67 accessions were ordered from IPK Gatersleben, Germany. In addition, three *V. faba* accessions were included, two with the wild type *VC1* allele and one homozygous for *vc1*. After an initial investigation, seeds of 23 accessions of 9 further species in subgenus *Vicia* were obtained from IPK.

### Plant growth and seed production

The seeds were inoculated with a mix of two *Rhizobium leguminosarum* bv *viciae* races and one *Bradyrhizobium* sp. (Elomestari OY, Tornio, Finland), to increase the probability that nodulation would take place. Each accession was sown into three 2-litre pots, filled with rough potting mixture (Kekkilän karkea ruukutusseos, Kekkilä Oy, Vantaa, Finland), with the aim of producing three plants per pot. Daylength in the glasshouse was set to 14 h light and 10 h dark, with 22°C during the day and 15°C at night. Growth continued for 11 months, during which the plants were given PK fertilizer twice.

Some species had not started flowering by four months, so two of the three pots were taken outside, covered with horticultural fleece and allowed to vernalize for two weeks. The average air temperature was 2.2°C but, inside the fleece, temperatures remained between 5° and 10°C. Seeds were gathered as they matured, the maturation period differing widely between the species. The earliest, *V. narbonensis* L. began maturing at four months, while the latest, *V. unijuga* A. Braun, commenced at 10 months. Eight of the 70 accessions did not produce any seeds. Seeds were also gathered from lentil (*Lens culinaris* Medik. (*Vicia lens* (L.) Coss. & Germ.)) and pea (*Pisum sativum* L. (*Lathyrus oleraceus* Lam.)) as an outgroup.

A new batch of seeds was sown after the VC measurements of the first seeds, comprising 23 new accessions, as well as the 11 that had produced few or no seeds earlier. The seeds were inoculated the same way as previously, then sown in small paper pots. After emergence, they were placed in a +4°C room for five weeks for vernalization. Three vigorous seedlings from each accession were transferred to 2-liter pots filled with the same potting mixture as previously, with three replicate pots of each accession. The pots were placed in a caged area in the University of Helsinki Viikki campus greenhouse for the summer growing season and were given PK fertilization twice during this time. Fourteen of the 34 accessions failed to produce any seeds. Leaves were collected from all individual pots before seed set for DNA extraction and analysis. The leaf samples were frozen and stored at -20°C until use.

### Analysis of VC content

A 1 g sample of seeds from each accession in the first set of species was milled using an MM300 TissueLyser Lab Vibration Mill Mixer (Retsch, Haan, Germany). A 0.5 g subsample was extracted with water (MilliQ), then vicine and convicine content measured on an HPLC platform (Waters Corporation, Milford, MA, USA) using an internal standard, as described earlier (Pulkkinen *et al*., 2015). For the second set of species, a Waters Acquity UHPLC system was used. The same protocol was used to process the samples as for HPLC, except that an Acrodisc 13 mm minispike with 0.2 µm wwPTFE membrane (Cytiva, Port Washington, NY, USA) was used before UHPLC analysis. UHPLC was run with the same setup as in (Wang *et al*., 2024), except that the water contained 0.5% formic acid, instead of 0.1%. The analytical performance for vicine was evaluated by establishing calibration curves. The limits of detection (LOD) and of quantitation (LOQ) for vicine were determined to be 0.006 mg/g and 0.018 mg/g, respectively, based on the standard deviation of the response and the slope of the calibration curve.

### DNA extraction and analysis

DNA was extracted from the leaves of the first batch of plants using the cetyltrimethylammonium bromide (CTAB) protocol (Doyle and Doyle, 1990) with the addition of 2.8 μl proteinase K (Thermo Fisher Scientific Baltics UAB, Vilnius, Lithuania) per 0.9 ml of CTAB.

PCR primers based on established *VC1* and *ribAB* sequences in *V. faba* were used (Björnsdotter *et al*., 2021). The forward primer was GGCAGCACAGTTGGCAACACC and reverse primer TGCAAGCTGATTTCCACAGTCAC. These fragments in faba bean span from exon 3 to exon 6, including introns 3, 4 and 5, and were the longest obtainable from all the species. The PCR reaction contained 1 μl (100 ng) of gDNA, 10 μM of forward and reverse primers, 2 μl Dreamtaq buffer, 0.2 μl DreamTaq DNA Polymerase EP0702 (Thermo Scientific, Waltham, MA, USA), μl dNTPs, and 14.4 μl of water (MilliQ) for a 20 μl reaction. The PCR program comprised: 1, 92°C for 120 sec; 2, 94°C 30 sec; 3, 54°C for 30 sec; 4, 72°C for 130 sec; steps 2-4 repeated 40 times; 5, 72°C 5 min. The products were separated on 1% agarose gels with 1 x TAE buffer at 120 V for 50 minutes. The fragments ranged in size from 1000 bp to 2200 bp and were cloned using pGEM-T Vector Systems I and JM109 competent cells (Promega, Madison, WI, USA). Three clones were sequenced from each accession. Plasmid DNA was extracted using GeneJET Plasmid Miniprep Kit (Thermo Fisher) and sent to Eurofins Genomics (Ebersberg, Germany) for sequencing.

Sequences from internal transcribed spacer II (*ITS2*) and mitochondrial cytochrome oxidase subunit I (*COX1*) were acquired from an MSc thesis produced on the same leaf samples as used in this study (Shehbala, 2021).

### Statistical and sequence analyses

The sequences were cleaned and assembled with ApE (Davis and Jorgensen, 2022). All sequences in a given species were aligned with Clustal Omega (Madeira *et al*., 2024) to check for anomalies. Sequences that did not have both primers were discarded; these discarded sequences were part of much larger *ribAB* sequences. Some species produced several bands, for example, *V. cracca* L. had three, at 1000 bp, 1400 bp and 1600 bp, which were from different *ribAB* forms. The sequence most similar to *VC1* was chosen from each species. A phylogenetic tree was constructed with MEGA12: Molecular Evolutionary Genetics Analysis version 12 (Kumar *et al*., 2024) with the maximum likelihood method and Kimura 2-parameter model.

Three sequences from faba bean cDNA using the same primers were included that code for known *ribAB* copies.

The ApE program was used to remove introns from the sequences so that an exon-only data set was produced. Both the exon-only dataset, as well as the one including introns, were used to determine nucleotide diversity using the DNAsp program (Rozas *et al*., 2017). Alignments for this were made with MEGA12. The dataset on VC content was subjected to one-way analysis of variance using SPSS 27 (IBM Corp., Armonk, NY, USA).

## Results

### Vicine-convicine occurrence

Of the 30 *Vicia* species measured for vicine and convicine, 14 had both compounds and 6 had vicine alone (Table 1). There was no species with convicine alone. All of the vicine-producing species were in the subgenus *Vicia*. Within the subgenus, the amount of vicine was relatively consistent within each section, with the highest amounts being in section *Vicia* and the lowest in section *Narbonensis*. No vicine or convicine was detected in the outgroup comprising lentil and pea. Within each species, accessions differed in VC content to a varying degree, with *V. angustifolia* L. showing the widest variation and *V. hyrcanica* Fisch. & C.A.Mey., with its very low total content, showing no detectable variation. *V. orobus* DC., *V. ochroleuca* Ten., and *V. amurensis* Oett. did not yield enough seed for VC analysis, so are missing from this table.

**Table 1.**
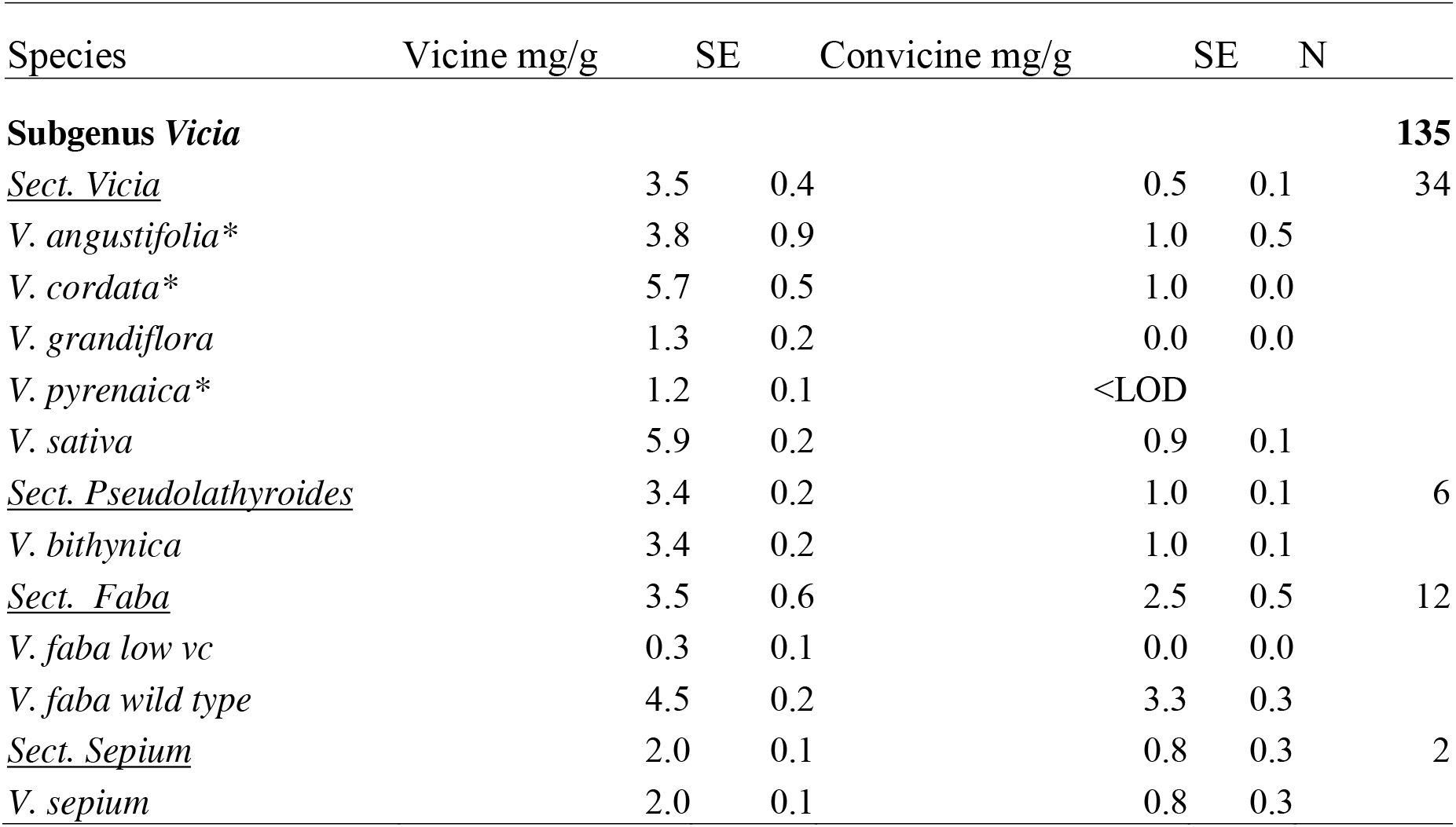
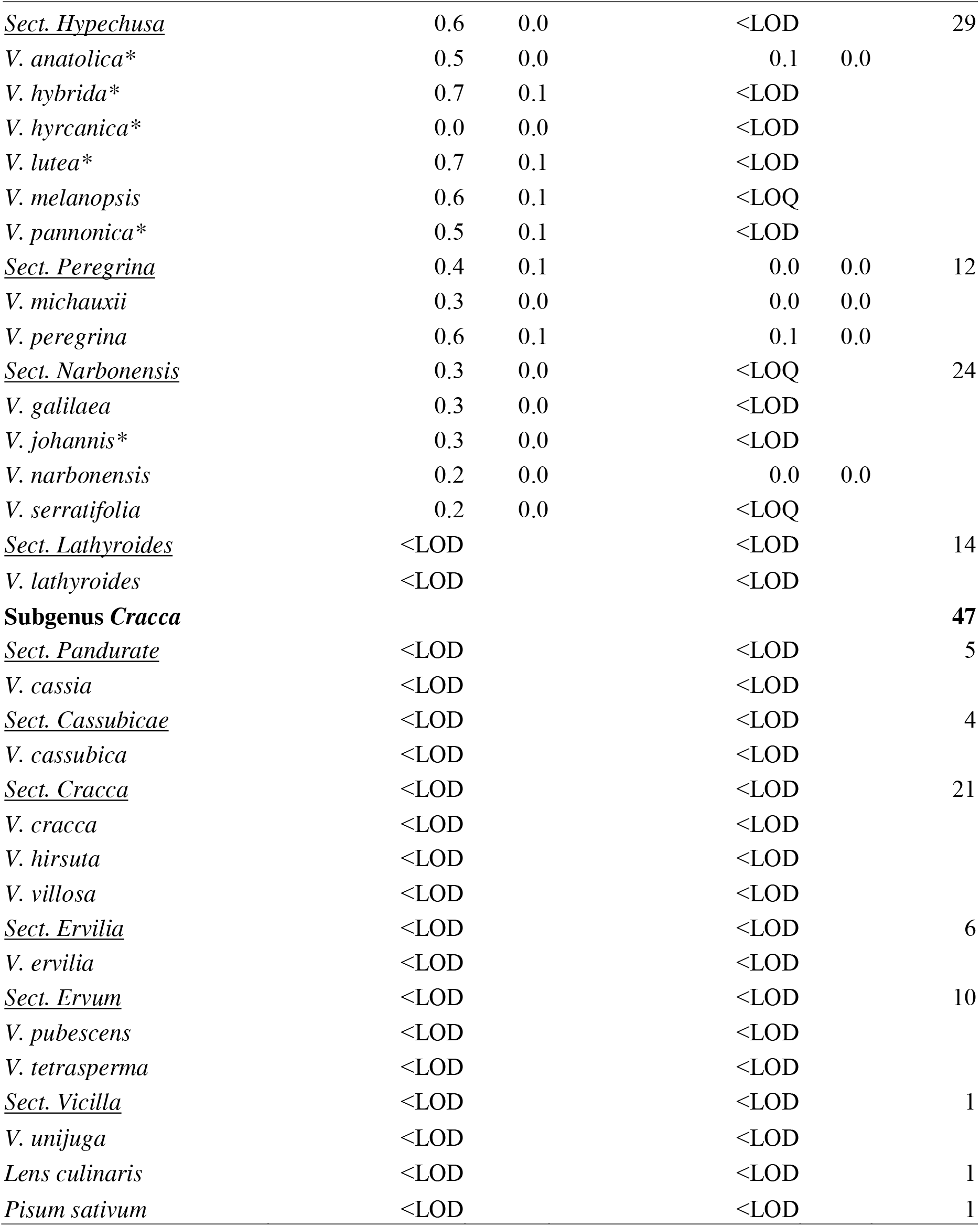
Vicine and convicine content of Vicia species and relatives, determined by HPLC or (*) UHPLC. Data show means and standard errors of two technical replicates of each accession and up to 3 accessions per species, giving a total of N samples.

The differences between sections in VC content were significant (*p* < 0.001). These sections also formed two groups, one (sections *Vicia, Faba, Pseudolathyroides*, and *Sepium*) where all the species produced over 1 mg/g of vicine, and the other (sections *Hypechusa, Peregrina*, and *Narbonensis*) with less than 1 mg/g. Convicine did not have similar groupings, with 6 species producing over 0.1 mg/g of convicine, 8 producing less than 0.1 mg/g, and the highest value being in section *Faba*.

### Presence of the VC1 gene

Following initial tests of several primer pairs, designed for the faba bean *VC1* sequence, on the original set of 67 accessions, the primer pair that worked on all accessions and produced the longest product was chosen. The amplification site started at *VC1* exon three and ended in exon six, spanning most of the coding region for RibB and half of the coding area of RibA. It includes the AT insertion site that is found in the recessive *vc1* allele.

The amplicons produced by the PCR were of consistent size within each subgenus, particularly in subgenus *Vicia* where they were around 1000 bp (Figure 1). Other amplicons were up to 2700 bp in size in *V. cassubica* (lane 25) and *V. orobus* (lane 26). Several species (*V. peregrina, V. lutea, V. orobus, V. cracca, V. villosa* ) produced more than one band.

**Figure 1.**
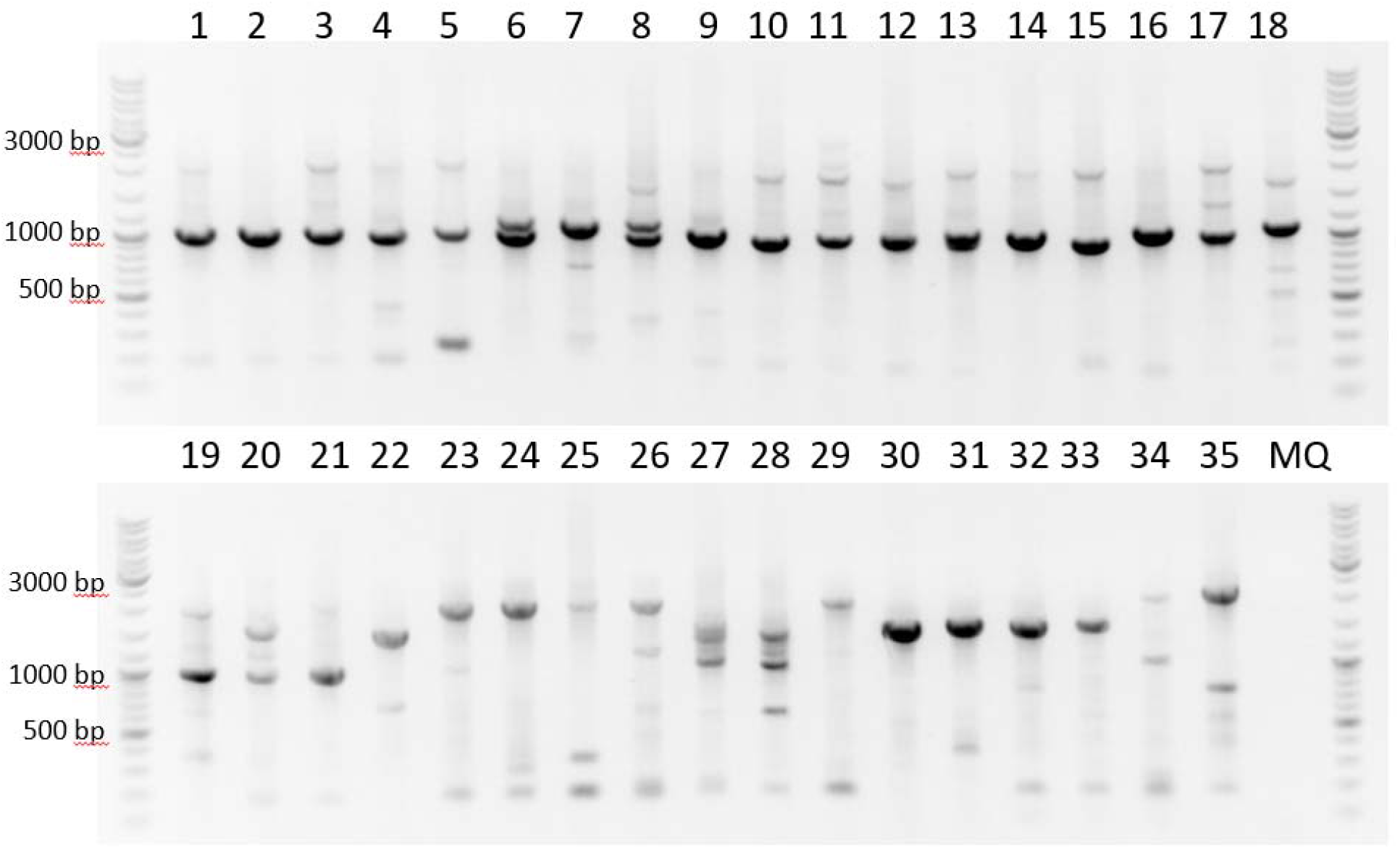
PCR amplification of a region of VC1. 1. V. sativa, **2**. V. cordata, **3**. V. angustifolia, **4**. V. grandiflora, **5**. V. sepium, **6**. V. peregrina, **7**. V. michauxii, **8**. V. lutea, **9**. V. hybrida, **10**. V. anatolica, **11**. V. melanopsis, **12**. V. pannonica, **13**. V. hyrcanica, **14**. V. pyrenaica, **15**. V. lathyroides, **16**. V. narbonensis, **17**. V. johannis, **18**. V. faba, **19**. V. galilaea, **20**. V. serratifolia, **21**. V. bithynica, **22**. V. hirsuta, **23**. V. amurensis, **24**. V. unijuga, **25**. V. cassubica, **26**. V. orobus, **27**. V. cracca, **28**. V. villosa, **29**. V. ochroleuca, **30**. V. cassia, **31**. V. ervilia, **32**. V. tetrasperma, **33**. V. pubescens, **34**. Lens culinaris, **35**. Pisum sativum. Lanes **1-21** belong to subgenus Vicia, **22-33** subgenus Cracca. ALT TEXT: Agarose gel electrophoresis image of PCR products from 35 samples. A DNA size marker is included for fragment size estimation. Lanes 1–21 show a strong band at approximately 1000 bp. Lanes 22–35 show larger fragments ranging from just over 1000 bp to approximately 2700 bp. The final lane contains a negative water control and shows no visible amplification product.

Phylogenetic analysis of the amplified sequences was applied to enable determination of the distribution of VC in the genus. The phylogenetic tree (Figure 2) shows that the sequence of the *VC1* fragment diverged from other *ribAB* fragments at a point corresponding to the node for subgenus *Vicia*, with the trait of producing VC being highly linked to the presence of *VC1*.

**2Figure 3.**
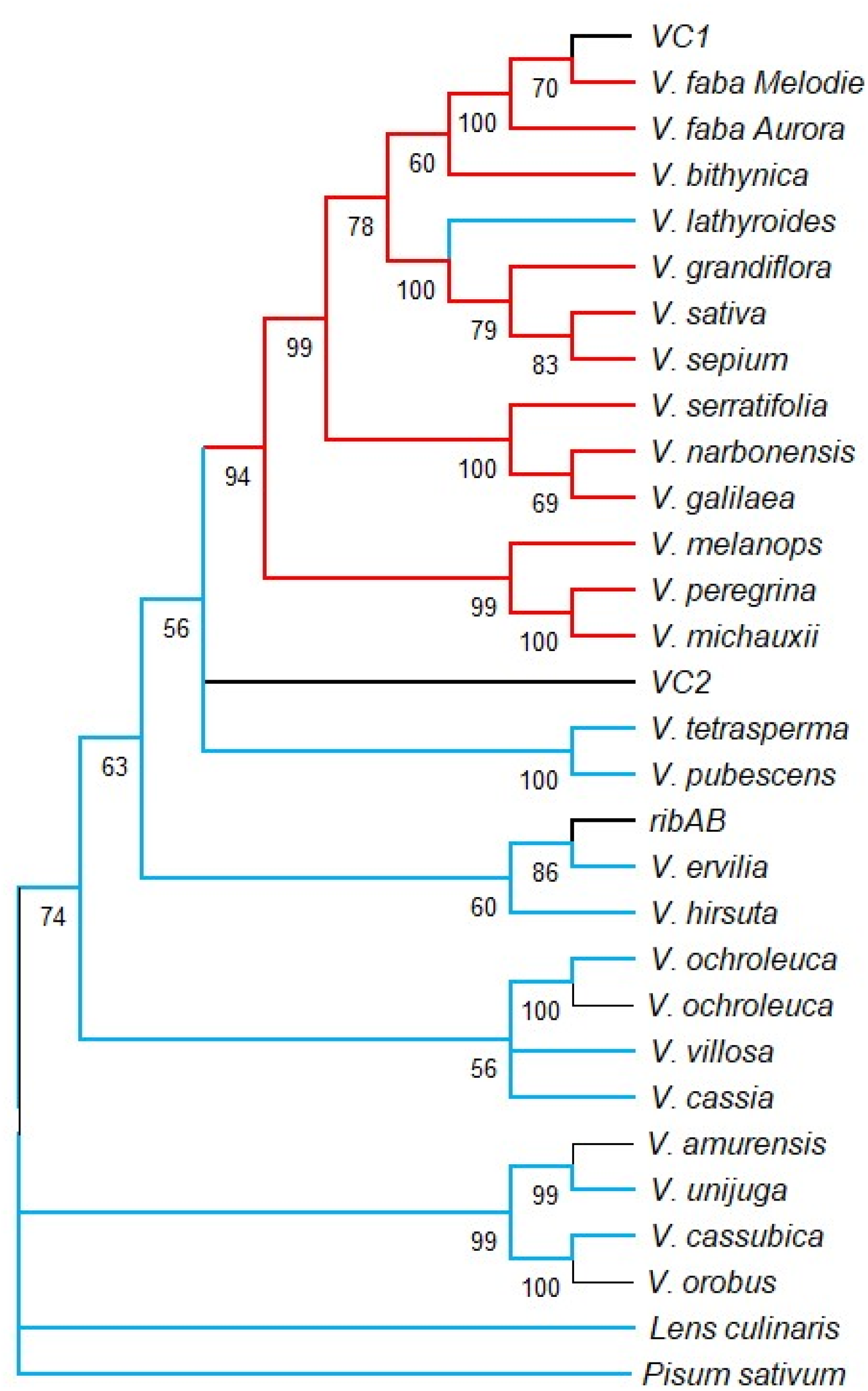
Phylogenetic tree of VC1 and ribAB sequences in genus Vicia along with lentil and field pea and the reference sequences of VC2 and seedcoat ribAB from V. faba line Hedin/2 (Jayakodi et al., 2023), assembled by the maximum likelihood method and the Kimura-2 model using MEGA12. Red color indicates the presence of vicine in the species. Blue lines indicate species that have no detectable vicine or convicine. Black that sufficient seeds were not obtained for VC analysis. ALT TEXT: Phylogenetic tree of *VC1* and *ribAB* sequences from species of the genus *Vicia*, together with lentil (*Lens culinaris*), field pea (*Pisum sativum*), and the reference sequences *VC1, VC2* and seed coat *ribAB* from Vicia faba line Hedin/2 (Jayakodi et al., 2023). The tree was constructed using the maximum-likelihood method and the Kimura 2-parameter model in MEGA12. Red labels indicate species in which vicine was detected, blue labels indicate species with no detectable vicine or convicine, and black labels indicate species for which insufficient seed material was available for vicine/convicine analysis. Notably, all vicine-containing species (red) cluster within a single branch of the tree, with *Vicia lathyroides* being the only non-vicine species (blue) within this clade.

Sequence diversity was substantially greater in *VC1* of subgenus *Vicia* than in *ribAB* of subgenus *Cracca* (Table 2). The diversity was much lower in the housekeeping genes *COX1* and *ITS2*. When the introns were excluded from the sequence analysis, *ribAB* sequence diversity was similar to that of *ITS2*. Non-synonymous mutations were six times greater in genomic sequences of *VC1* of subgenus *Vicia* than in *ribAB* of subgenus *Cracca* (Table 3). Synonymous mutations, however, were at comparable rates in both subgenera.

**Table 2.** Diversity in 67 genomic or exon sequences of VC1 in subgenus Vicia and RibAB in subgenus Cracca, as determined by analysis with DNAsp, along with diversity in 24 sequences of housekeeping genes COX1 and ITS2. Pea and lentil were not included in the analyses.

| Subgenus | N = 67 |  | N = 24 |  |  |  |
| --- | --- | --- | --- | --- | --- | --- |
|  | VC1 / *ribAB* |  | VC1 / *ribAB* |  | COX1 | ITS2 |
|  | Genomic | Exon | Genomic | Exon | Genomic | Genomic |
| **All** | 0.069 | 0.064 | 0.088 | 0.064 | 0.003 | 0.027 |
| ***Vicia*** | 0.067 | 0.061 | 0.106 | 0.062 | 0.003 | 0.025 |
| ***Cracca*** | 0.064 | 0.035 | 0.097 | 0.037 | 0.002 | 0.022 |

**Table 3.** Proportion of synonymous mutations (S), non-synonymous mutations (N) and their ratio in the genus Vicia as a whole, subgenus Vicia and subgenus Cracca, following analysis with DNAsp. Pea and lentil were not included in the analyses.

| Subgenus | N | Pi(s) [dS] | Pi(a) [dN] | dN/dS |
| --- | --- | --- | --- | --- |
| All | 67 | 0.146 | 0.038 | 0.260 |
| *Vicia* | 35 | 0.115 | 0.043 | 0.375 |
| *Cracca* | 32 | 0.121 | 0.007 | 0.061 |

## Discussion

This work has shown that VC synthesis is found only in subgenus *Vicia* and is closely associated with the *VC1* variant of *ribAB*. Additional *ribAB sequences* from *V. faba* clustered with the *ribAB variants* found in subgenus *Cracca*. One species in the subgenus *Vicia* had a *VC1* sequence that was closely related to the other *VC1* sequences, but its VC content was below the limit of detection. The results are compatible with a monophyletic origin of *VC1* and of VC synthesis in subgenus *Vicia*, but do not provide information on the phylogeny of the rest of the genus.

VC production is present in all sections of subgenus *Vicia*, pointing to its appearance at the evolutionary point when subgenera *Vicia* and *Cracca* separated from each (Schaefer *et al*., 2012). The trait accompanies the identifiable *VC1* sequence, which differs from other *ribAB* copies (Björnsdotter *et al*., 2021; Jayakodi *et al*., 2023). *VC1* is consistent within the subgenus *Vicia* and it differs from the two other *ribAB* forms found in *V. faba*.

The notable exception is *V. lathyroides* L., which apparently has the *VC1* gene but, unlike every other tested species in subgenus *Vicia*, does not produce VC. The placement of *V. lathyroides* in the subgenus *Vicia* is otherwise solid and confirmed by other (Hanelt and Mettin, 1989; Leht, 2009). Work is currently under way on identifying how *V. lathyroides* has silenced its VC production.

Convicine was produced by only those species where vicine production exceeded 1 mg/g by the final dry weight of the seed, and was present in all species where this value exceeded 2 mg/g. Similarly, faba bean cultivars with low VC content due to homozygosity of the *vc1* allele have no detectable convicine (Khamassi *et al*., 2013). These observations show that convicine is a secondary product after vicine. They are in line with the current model of VC biosynthesis, wherein a deamination takes place after the initial action of *VC1* to produce DARPP (Björnsdotter *et al*., 2021). Factors affecting the relative amount of convicine have not been identified; its content exceeds that of vicine in only a few faba bean genotypes (Khamassi *et al*., 2013).

Within the subgenus *Vicia*, the sequence-based phylogeny of *VC1* (Figure 2) agrees with the morphology-based phylogeny of Leht (Leht, 2009). Elsewhere, the sequence-based phylogeny based on *ribAB* is less consistently informative. Some pairs of species in the same section, such as *V. unijuga* and *V. amurensis*, as well as *V. tetrasperma* and *V. pubescens*, cluster with each other, conforming with the morphology-based phylogeny. The two *ribAB* forms obtained from *V. faba* are closely related to the rest of the *ribAB* sequences, demonstrating the differences between *VC1* and other *ribAB* forms. Field pea and lentil behaved as out-groups, as expected.

*ITS2* and *COX1* were included to allow comparison of sequence diversity in other genes in these subgenera with that of *ribAB* and *VC1. COX1* diversity was very low, at 0.002, consistent with the slow rate of mitochondrial gene evolution in plants (Knoop, 2004). *ITS2* diversity was ten times higher, in line with its faster evolution (Poczai and Hyvönen, 2010). Sequence diversity in *ITS2* of subgenus *Cracca* was lower than that in subgenus *Vicia*, despite subgenus *Cracca* being the larger group with notoriously hard to define taxonomic borders (Kupicha, 1976; Hanelt and Mettin, 1989).

The difference between dN/dS ratios in genomic and exon-only sequences of *ribAB* shows that most of the non-synonymous mutations were in the introns. The exon sequences were nearly as conserved as the genomic sequences of *ITS2*, demonstrating the purifying selection on this gene and the essential role of riboflavin synthesis in the tissue in which *RibAB* is expressed (Hiltunen *et al*., 2012). The dN/dS ratios were much lower in r*ibAB* than in the *VC1* exon sequences. This result indicates that *VC1* is not under stabilizing or purifying selection to the same extent as *ribAB* and is consistent with the hypothesis that the gene is evolutionarily a relatively recent and accessory introduction. Gene duplication resulting in novel molecular functions are common in plants and have resulted in many traits in domesticated plants (Panchy *et al*., 2016). Examples of this include gene duplication leading to capsaicin biosynthesis in *Capsicum* genus (Qin *et al*., 2014) and diversity in fruit shape in domesticated tomato (Xiao *et al*., 2008).

In conclusion, this study supports a monophyletic origin of VC biosynthesis within subgenus *Vicia*, and a strong association of the trait with the presence of the *VC1* form of *ribAB. V. lathyroides* is an important exception, as it carries *VC1*, but produces no VC. The presence of convicine depends on vicine, which supports the proposed VC biosynthetic pathway, in which convicine is a secondary product derived from the vicine pathway. Finally, the increase in sequence diversity between *VC1* and *ribAB* indicates different evolutionary constraints, with strong purifying selection removing non-synonymous substitutions from *ribAB* forms. Together, these findings expand our understanding of VC biosynthesis evolution and provide information for future studies within the genus.

## Abbreviations

VC: Vicine and convicine
DARPP: 2,5-diamino-6-ribosylamino-4(3H)-pyrimidinone 5’-phosphate

## Supplementary data

1. Accession numbers of the plant material used in this study
2. DNA sequences

## Acknowledgements

The authors gratefully acknowledge the German Federal Ex Situ Genebank at the Leibniz Institute of Plant Genetics and Crop Plant Research (IPK), Gatersleben, Germany, for providing the accessions used in this study. We also thank the Genebank Information System (GBIS) team for facilitating accession selection and access to associated passport data.

We thank Sanna Peltola, Marjo Kilpinen, Anne-Marie Narvanto, and Eyerusalem Shehbala for their valuable technical assistance and professional support throughout this study.

## Author contributions

Conceptualization: LLV, AHS, FLS; Methodology and investigation: all authors; Writing – original draft: LLV, FLS; Writing – review & editing: LLV, AHS, FLS; Supervision: FLS, AHS; Funding acquisition: LLV, FLS. The final submission was approved by all authors.

## Conflict of interest

The authors declare that they have no known competing financial interests or personal relationships that could have appeared to influence the work reported in this paper.

## Funding

LLV was supported by Agronomiliitto through the Suomi Kasvaa Ruuasta programme (grant numbers 20190010, 20200111, and 20210084).

## Data availability

The data supporting the findings of this study are available within the article and its supplementary material.

## Notes

### Competing Interest Statement

The authors have declared no competing interest.

